# Uncertainty avoidance versus conditioned reinforcement: exploring paradoxical choice in rats

**DOI:** 10.1101/2021.08.12.456071

**Authors:** Victor Ajuwon, Andrés Ojeda, Robin A. Murphy, Tiago Monteiro, Alex Kacelnik

## Abstract

Paying a cost to reduce uncertainty can be adaptive, because better informed decision-makers can align their preferences to opportunities. However, birds and mammals display an appetite for information that they cannot use to functionally alter behaviour or its outcomes. We explore two putative motivational mechanisms for this paradoxical behaviour. The ‘information hypothesis’, proposes that reducing uncertainty is reinforcing *per se,* consistent with the concept of curiosity: a motivation to know, in the absence of instrumental benefits. In contrast, the ‘conditioned reinforcement hypothesis’ sees information-seeking as a consequence of asymmetries in secondarily acquired reinforcement: responding increments caused by post-choice stimuli announcing positive outcomes (S^+^) exceed decrements caused by stimuli signalling absence of reward (S^−^). We contrast these hypotheses experimentally. Rats chose between two equally profitable options delivering food probabilistically after a fixed delay. In the informative option (*Info)*, the outcome (food/no food) was signalled immediately after choice, whereas in the non-informative option (*NoInf*o) outcomes were uncertain until the delay lapsed. Subjects preferred *Info* when (1) outcomes were signalled by salient auditory cues, (2) only the absence of reward was signalled, and (3) only reward was signalled, though acquisition was slower when rewards were not explicitly signalled. Our results show that a salient good news signal is not required as a conditioned reinforcer to generate paradoxical preferences. Terminal preferences support the information hypothesis but the slower acquisition of *Info* preference when S^+^ is not present is consistent with the conditioning account. We conclude that both uncertainty reduction and conditioned reinforcement influence choice.

## Introduction

Models of instrumental learning argue that animals increase the frequency of actions that result in higher probability of desirable consequences such as food or water, and reduce the frequency of actions that enhance undesirable ones, such as food absence or other aversive consequences, with desirability and aversiveness assumed to have adaptive roots (for a review see Staddon and Cerutti, 2003). Similar algorithms are used in artificial reinforcement learning processes (Sutton and Barto, 2018). Essential commodities or other substantial beneficial outcomes serve as positive reinforcers, and their absence as negative ones, and it is not surprising that in the laboratory such events modulate animals’ lever pressing or key pecking responses. However, a question that has recently seen a resurgence of interest in the psychological (Cunningham and Shahan, 2018; Shahan and Cunningham, 2015), neuroscientific/robotics (Gottlieb and Oudeyer, 2018; van Lieshout et al., 2020), and computational (Dubey and Griffiths, 2020) literatures, is whether information *per se* (reductions in uncertainty) can modulate behaviour through the same learning processes that conventional rewards do, that is, whether information can act as a primary reinforcer.

In a world where uncertainty is pervasive, information is a valuable asset that can be used by decision-makers to enhance efficiency in activities such as foraging, to improve performance (Behrens et al., 2007; Dall et al., 2005). In experimental instrumental tasks, animals may seek information before making choices (Gottlieb et al., 2014) and this can improve the acquisition of appetitive commodities available to them (Foley et al., 2017; Kobayashi and Hsu, 2019). In this context, the adaptive value of information-seeking derives from its ability to help increment some well-defined benefit. What appears paradoxical is that animals value information even in cases where it has no potential instrumental use - that is, they seek out information ‘for its own sake’, are ‘uncertainty averse’, or are ‘curious’ (Kidd & Hayden, 2015; Bromberg-Martin and Hikosaka 2009; Cervera et al., 2020)

The idea that individuals find information intrinsically rewarding has been suggested as an explanation for ‘observing response’ experiments, first carried out by Wyckoff (1951, *unpublished thesis;* see Wyckoff, 1969). In this paradigm, subjects can seek information about forthcoming contingencies by performing a response that triggers reward-predictive signals, though the information cannot be used to modify outcomes. Wyckoff presented pigeons with a white key and two types of trials. In rewarded trials pecking the key resulted in food delivery after 30s (a fixed-interval 30 schedule; FI30), while in unrewarded trials pecking at the key did not produce food (an extinction schedule; Ext). The system alternated periodically between FI30 and Ext. The critical aspect was the addition of a pedal such that if the animal stepped on it, then the white key turned red during the rewarded trials and green during the unrewarded trials. Hence, red was a positive stimulus (S^+^) anticipating food while green was a negative stimulus (S^−^) signalling no food. In other words, the pedal response informed the animal of the current state of the world, but did not modify it. Stepping on the pedal was labelled the ‘observing response’. The pigeons readily acquired and maintained the observing response even if this did not change the availability of food. Similar procedures have been conducted with variable delays to food (Bower et al., 1966) and aversive outcomes such as electric shocks (Lockard, 1963). In all cases animals display the ‘observing response’; they choose to elicit signals that resolve uncertainty about probabilistic future outcomes.

The observing response is intriguing because the information provided by the signals cannot be used by subjects to modify or enhance any substantive adaptive outcome, so it is not clear why animals value them, or how the response is acquired by reinforcement learning. Also, the result appears to contradict normative models of reward-maximisation in other fields (e.g., Mas-Colell et al., 1995; Stephens and Krebs, 1986) that do not predict actions that result in functionless information. A number of theoretical hypotheses have been proposed to explain observing responses (for a review see Dinsmoor,1983). One possible mechanistic explanation, the “information hypothesis”, suggests that animals find information itself intrinsically rewarding because it relieves uncertainty, which is aversive (Berlyne, 1960, 1957; Hendry, 1969). According to this account, information is a primary reinforcer that can modulate behaviour. Functionally, this could emerge evolutionarily if information is generally associated with substantive benefits in ecological contexts. This view is consistent with notions of ‘curiosity’ defined as the motivation to ‘know’ for the sake of it, or acquire information in the absence of instrumental incentives (Cervera et al., 2020; Gottlieb and Oudeyer, 2018; Kidd and Hayden, 2015). The idea that individuals value information has also recently been explored in humans. Bennet et al. (2016) suggested that information may be valued because it prevents temporally-prolonged uncertainty. Other investigators have proposed that information may derive intrinsic reinforcing value through reward prediction errors that enable subjects to appetitively savour good news about positive outcomes (Brydevall et al., 2018; Iigaya et al., 2016).

An alternative mechanistic explanation, the ‘conditioned reinforcement hypothesis’ (Bower et al., 1966; Prokasy, 1956; Wyckoff, 1959), proposes that the signal for food (S^+^) in observing response tasks acquires secondary reinforcing properties and becomes a conditioned reinforcer. By definition, a reinforcer is an event that modifies the frequency of a response when the event is contingent on that response. For example, the presentation of food is a positive excitatory reinforcer because when it is contingent on a lever being pressed, animals will press the lever more frequently than when the lever pressing is not paired with food. Conditioned reinforcers are initially neutral stimuli that themselves become reinforcing after having been paired with a primary reinforcer (see Mackintosh, 1974, and for applications in machine learning see Sutton and Barto, 2018). Thus, it has been argued that the subsequent production of S^+^, once it has been associated with food, is what drives animals to respond in observing response and similar experiments. However, by the same reasoning, an S^−^ predicts a reduction in food probability, and might be expected to acquire the power to reduce the frequency of responding (i.e., to acquire aversive, or inhibitory properties). If these two effects are of different absolute magnitude (favouring S^+^), conditioned reinforcement offers a descriptive account of the acquisition of observing responses which is not dependent on the animal being sensitive to uncertainty or its reduction.

Both the information and conditioned reinforcement hypotheses are plausible and distinct. From an animal learning perspective, the former proposes that information is a primary reinforcer, while the latter relies on the well-established phenomenon of conditioned reinforcement as an explanation. Though the information hypothesis is simple, functionally appealing, and intuitive, it fell broadly out of favour when evidence in pigeons began to emerge that was interpreted to be incongruent with it, but consistent with the conditioned reinforcement account (Dinsmoor, 1983; Roper and Zentall, 1999; Shahan and Cunningham, 2015).

The information hypothesis predicts that both S^+^ and S^−^ individually, should be sufficient to reinforce observing responses, because both provide information about an otherwise uncertain outcome. The conditioned reinforcement hypothesis, on the other hand, stipulates that only S^+^ should be positively reinforcing, thus being responsible for the acquisition and maintenance of observing responses. According to this view, although S^−^ reduces uncertainty just as much as S^+^, its presence should reduce rather than increase the observing response. To arbitrate between both hypotheses, researchers have carried out cue manipulation experiments in which either the good (e.g. food) or bad (e.g. no food) outcome is no longer preceded by a signal, or in other words the presentation of either S^+^ or S^−^ is omitted (Dinsmoor, 1983; Dinsmoor et al., 1972; Silberberg and Fantino, 2010). If information value is the driver of observing responses then the behaviour should be acquired and maintained by either S^+^ or S^−^ respectively, but if conditioned reinforcement is the cause, observing responses will not be acquired and/or maintained when the response generates a signal for bad outcomes (S^−^) but not good ones (S^+^). These cue manipulation experiments found that in pigeons, S^−^ alone was not sufficient to maintain observing, leading to the interpretation that information gain is not a sufficient incentive to generate preference (e.g., Dinsmoor et al., 1972; Jenkins and Boakes, 1973; Kendall, 1973; Silberberg and Fantino, 2010)

Recently however, results from novel protocols have rekindled interest in the possibility that animals find information intrinsically rewarding. Experiments in monkeys have found that they prefer to receive unambiguous signals about the magnitude of upcoming water rewards, over ambiguous or delayed signals, and are willing to forfeit water to do so. Furthermore, these preferences are correlated with activity in neurons implicated in the representation of primary rewards (Blanchard et al., 2015; Bromberg-Martin and Hikosaka, 2011, 2009). However, the psychological mechanism underlying these preferences is not clear. Daddaoua et al. (2016) showed that monkeys learn to actively search for positive Pavlovian cues to reduce uncertainty and obtain conditioned reinforcement, but the relative reinforcement provided by both of these factors is debatable. Taken together, currently available results show that the old conundrum of whether a reduction of uncertainty can on itself significantly reinforce behaviour is still unresolved.

Related experiments called ‘paradoxical’ or ‘suboptimal’ choice – similar in rationale to, and derived from, the observing response protocol - have found that pigeons (e.g., Fortes et al., 2016; González et al., 2020; Macías et al., 2021; McDevitt et al., 2018; Smith et al., 2016 also see McDevitt et al., 2016 and Zentall, 2016 for reviews), starlings (Vasconcelos et al., 2015) and rats (Cunningham and Shahan, 2019; Ojeda et al., 2018), prefer an alternative that provides information that they cannot use, not just when the information is neutral with respect to reward maximisation, but even when the informative option provides less reward. In this paradigm, both alternatives result in probabilistic food delivery after a delay. In the informative option, signals (S^+^ or S^−^) anticipate the trial’s outcome immediately after a choice response, while in the non-informative option subjects are uncertain about outcomes throughout the delay. Remarkably, pigeons and starlings choose the informative option when it gives 80% less reward than the non-informative alternative (Vasconcelos et al., 2015; Fortes et al., 2016), while rats can sacrifice at least 20% of potential rewards (Ojeda et al. 2018, Cunningham and Shahan, 2019) by selecting the informative option. The fact that animals forfeit potential food rewards to generate apparently useless signals is a strong reason to suspect a hypothetical primary reinforcing value of uncertainty reduction in observing response and paradoxical choice experiments.

To investigate whether uncertainty reduction or conditioned reinforcement can account for information-seeking behaviour in rats (*Rattus norvegicus)*, we conducted paradoxical choice experiments using cue manipulations involving the omission of S^+^ or S^−^ from the informative option. While manipulations to the signalling properties of choice alternatives have been performed in birds (e.g., Fortes et al., 2017; Vasconcelos et al., 2015 and see McDevitt et al., 1997 for a similar task), there are no reports of the relative quantitative impact of symmetrical omissions of S^+^ and S^−^, the most fundamental distinctive prediction of the two hypotheses. Thus, our experiment offers novel insights into the mechanisms underlying paradoxical choice behaviour in rats and potentially other species. Our results indicate that both uncertainty reduction and conditioned reinforcement underly information-seeking behaviour.

## Methods

### Experimental strategy and its rationale

In three treatments, subjects were exposed to repeated choices between two options of equal average profitability. Each option delivered reward with 50% probability, a fixed interval after being chosen. In all treatments, the two options only differed in their signalling properties (but see differences between treatments below): in the informative option (*Info*), the outcome of trials (food/no food) was signalled (or otherwise predictable) between each choice and the outcome, while in the other option (*NoInfo)* the outcome remained uncertain until it was realised. Because the signalling occurred post-choice in *Info*, it could not be used to modify the probability of receiving food.

The treatments differed in the signalling used in the *Info* option, as follows. In the *S*^+^_*S*^−^ treatment, the interval between choosing *Info* and the outcome was filled in rewarded or unrewarded trials by either of two sounds, namely S^+^ or S^−^ respectively. In the *Only_S*^−^ treatment, the interval was silent in trials when food was due, but filled with a sound when no food was coming. In the *Only_S*^+^ treatment, the same interval was filled with a sound signal in trials when food was forthcoming and with silence when it was not.

With experience animals in all treatments can infer when a trial will end in reward or not, but in the *Only_S*^−^ treatment food is not anticipated by a defined perceptual cue acting as predictor. The information hypothesis predicts acquisition and persistence of a preference for the *Info* option in all groups, because information is provided either by an acoustic signal or by its absence. However, assuming that only salient stimuli acquire secondary reinforcing properties, the conditioned reinforcement hypothesis predicts a preference for *Info* only when S^+^ is present, namely in the *S*^+^_*S*^−^ and *Only_S*^+^ groups, with the latter eliciting strongest *Info* preference. In the *Only_S*^−^ treatment responding to *Info* causes either no physical signal (when food is due) or a signal for bad news, hence precluding a simple secondary reinforcement account for *Info* preference. We recorded two measures of preference, namely proportion of choices in 2-option choice trials, and response latency (reaction time) in 1-option forced trials. The latter has proven to be a robust metric of preference in a variety of different behavioural protocols and species (viz. Kacelnik et al., 2011; Monteiro et al., 2020; Reboreda and Kacelnik, 1991; Sasaki et al., 2018; Smith et al., 2018).

### Subjects

All experiments were carried out in compliance with the UK Animal (Scientific Procedures) Act (1986) and its associated guidelines. 24 male Lister Hooded rats (provider Envigo), 11 weeks old at the start of the experiment served as subjects. Animals were housed in groups of four. Throughout the experiment they were food deprived to 85-90% of their free-feeding weight using growth curves from the provider. Initial weight: 337 ± 14, final weight: 357 ± 16 (mean ± std.) Water was provided ad-libitum in their home cages and they were maintained in a 12-hour dark/light cycle with lights on at 6 AM.

### Apparatus

Testing was carried out in eight operant chambers (Med Associates, USA.) Each chamber contained three retractable levers: one in the back panel (centre) and two in the front panel, left and right of a central food magazine delivery tray equipped with an infrared beam and a sensor to record head entry. Each reward delivery consisted of four 45mg sucrose pellets (TestDiet, USA). A speaker was positioned above the food magazine in the front panel. Each chamber was also equipped with a house light (white) and a fan, which were switched on for the duration of the session. The chambers were controlled via custom-written Med-State Notation programs running on MED-PC V (Med Associates, USA).

### Experimental procedures

#### General procedure

We used a trial-based chain procedure as displayed in Fig. 1. There were two kinds of trials: 2-option choice trials and 1-option forced trials. A day’s session was composed of 60 trials: 40 forced (half *Info* and half *NoInfo*) and 20 choice, which were randomly intermixed. All trials started with the rear lever (initial link) extending. Pressing this lever resulted in its retraction, and either one (forced trials) or both (choice trials) of the front levers being presented. Pressing a front lever could initiate an acoustic cue (the terminal link) and the retraction of that lever (forced trials) or both levers (choice trials). The auditory cue, if present, was broadcast for a 10 second interval, after which food delivery occurred in rewarded trials without the need for a further response. Thus, each option was programmed as a discrete trial, response initiated, fixed time 10s, partial reinforcement 50% schedule. Trials were separated by an inter-trial interval (ITI) generated by sampling from of a truncated Poisson distribution with a mean of 50 seconds (range: 10 – 120 seconds) and adding 10 seconds. A session finished after 60 trials or 3 hours, whichever occurred first.

**Fig. 1.**
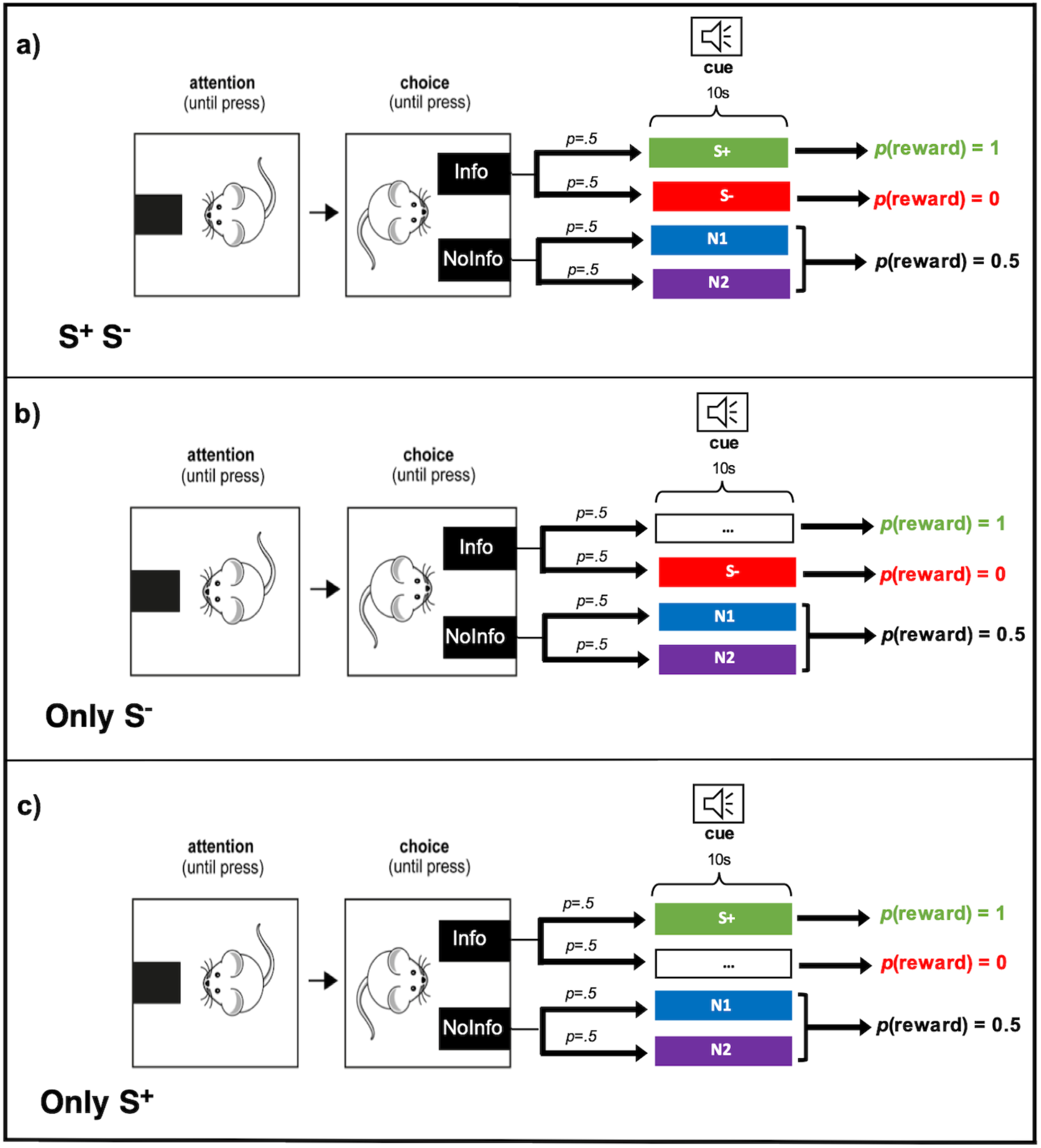
Experimental design showing choice trial structure for each treatment. Blank boxes with ellipsis indicate no auditory signal (silence) preceding outcomes. *p* denotes probability.

#### Experimental Procedure

In the *S*^+^_*S*^−^ treatment, choosing *Info* resulted with equal probability in either S^+^, which was paired with reward, or S^−^, which was paired with no reward, thereby reliably informing the subject of the forthcoming outcome. Responding to *NoInfo*, on the other hand, resulted with equal probability in either one of two cues: N1 or N2, which were both associated with a 50% probability of either outcome; therefore neither cue informed the subject of forthcoming reward.

The other two treatments, *Only_S*^+^ and *Only_S*^−^, were variations on the *S*^+^_*S*^−^ treatment, differing only in the signalling properties of *Info*. In *Only_S*^−^ responding to *Info*, resulted with equal probability (50%) in either a 10 second silence, followed by the delayed reward (the omission of a cue associated with reward i.e., S^+^), or the auditory S^−^ cue, which was associated with no reward. In *Only_S*^+^ choosing *Info* resulted with equal probability (50%) in either the cue *S*^+^, which was associated with reward after 10 seconds, or a 10 second silence (omission of the S^−^ cue) followed by no reward.

A between-subjects design was used, with eight rats in each group. Subject assignment to group was organised such that there was no correlation between group and any of the following parameters: side of the informative option; hour of testing; cage in which the animals were housed, or cue-reward contingencies. For each group, the subjects performed one daily session for 14 days. Each rat was trained at the same time every day; one cohort of rats began the experiment at 9:00 A.M., another at 12:30 P.M., and the last at 3:30 P.M.

#### Cues

The four cues consisted of sounds, all with a duration of 10 seconds, and each associated to a reward probability. There were two cues for the informative option: S^+^ (100% reward probability) and S^−^ (0% reward probability), and two cues for the non- informative option: N1 and N2 (both with 50% reward probability). The four sounds were: a low frequency pure tone (3 kHz, 78 dB), a high frequency pure tone (6 kHz, 78 dB), a buzzing sound (78 dB) and a clicking sound (74 dB). Assignment of sounds to reward probabilities was counterbalanced across subjects to avoid the possibility of option preferences being generated by any intrinsic aversive or attractive properties of the sounds.

### Training

#### Magazine training

To habituate the rats to the box and the delivery of food rewards, training began with a single variable interval session where food was delivered on average once a minute (VI60 free food schedule) a total of 60 times. The variable interval was sampled from a truncated Poisson distribution with a mean of 60 seconds and range of 0-120 seconds.

#### Lever training

Over the next three sessions the rats were trained to press the two front levers. Either lever (left or right with equal probability) was available on each trial (60 trials per session). Once a lever extended into the chamber, a single press resulted in its retraction and immediate reward delivery (Fixed Ratio 1 schedule). One of the levers then again became available after a delay composed of a constant duration plus a variable one. The constant component was 5 seconds, and the variable one was sampled from a truncated Poisson distribution with a mean of 20 seconds and a range of 5 – 60 seconds. All three sessions concluded after 60 reward deliveries i.e. 30 lever presses on each side, or after 3 hours.

#### Cue training

To train the subjects to the reward contingencies of the four auditory cues (S^+^, S^−^, N1, and N2), the main experiment was preceded by a Pavlovian protocol in which all the rats were exposed to the cues and their respective reward contingencies. In this phase, cue presentation was independent of the behaviour of the rat. These cue-training sessions consisted of 40 trials, with 10 trials for each cue randomly intermixed. To avoid large deviations from the expected outcome probabilities of cues N1 and N2 in each session, proportions of outcomes were fixed as one half for each cue. Trials were separated by an ITI generated by sampling from of a truncated Poisson distribution with a mean of 50 seconds (range: 10-120 seconds) + 10 seconds. Subjects performed one daily session of this phase for 10 days. Cumulative time spent head-poking into the food magazine was measured to establish the degree of cue discrimination.

### Data Analysis

Data processing and analysis was carried out in MATLAB 2017a and statistical tests were carried out with R statistical software (Version 1.2.5033). A type-1 error rate of 0.05 was adopted for all statistical comparisons and the Tukey test was used for all multiple comparisons. For statistical analysis, proportion data was arc-sine square-root transformed to normalize the residuals. Head-poking data as well as latency index data, were square root transformed (Grafen and Hails, 2002).

Mean choice proportion data for each treatment group were fitted with sigmoidal curves using the following function:

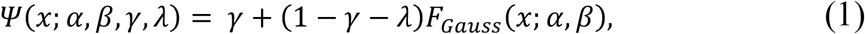

with *F*_*Gauss*_(*x*; *α*, *β*) a cumulative Gaussian function (Fig. 3). Non-linear least squares was used and implemented with the FitPsycheCurveWH function in MATLAB (Wichmann and Hill, 2001). λ and *γ* set the upper and lower bounds of the curves respectively while *α* gives the inflection point and *β* the slope. The upper bound was set at 1 for all curves while other parameters were estimated (Table S1).

To measure preference on the basis of latency to respond in forced trials, for each individual we calculated an index, *L*_*(Info)*_, using the median latencies to respond on *Info* (*R*_*(Info)*_) and *NoInfo* (*R*_*(NoInfo)*_) forced trials for each session: *L*_*(Info)*_ = *R*_*(Info)*_ / (*R*_*(Info)*_ + *R*_*(NoInfo)*_). Values of *L*_*(Info)*_ < 0.5 or *L*_*(Info)*_ > 0.5 indicate a preference for *Info* or *NoInfo* respectively, as measured in forced trials, independently of the measure of preference based on choices in 2-option trials.

## Results

### Training

#### Cue discrimination

A condition for the interpretation of preferences is that subjects were able to discriminate the contingencies of each cue; we examined this using cumulative head-poking time during the 10s interval between choice and outcome, when the cues were present, pooling data from the last three training sessions, across the groups, which up to that point had no differential experience (Fig. 2). The cue associated with 100% reward probability (S^+^) had the longest cumulative head poking duration (2.98s ± 0.14; mean± s.e.m.), followed by the average of both cues associated with 50% probability (N1 & N2: 2.40s ± 0.13), while the cue associated with no reward (S^−^) elicited the shortest average head poking duration (1.19s ± 0.09). A one-way repeated-measures ANOVA revealed a significant effect of cue (F_2,46_ = 44.38, *P* < 0.0001). Post-hoc pair-wise comparisons showed a significant difference between all pairs (100% vs 50%, *P* < 0.05; 100% vs 0%, *P* < 0.001 and 50% vs 0%, *P* < 0.001). This confirms that subjects discriminated the contingencies programmed for each cue.

**Fig. 2.**
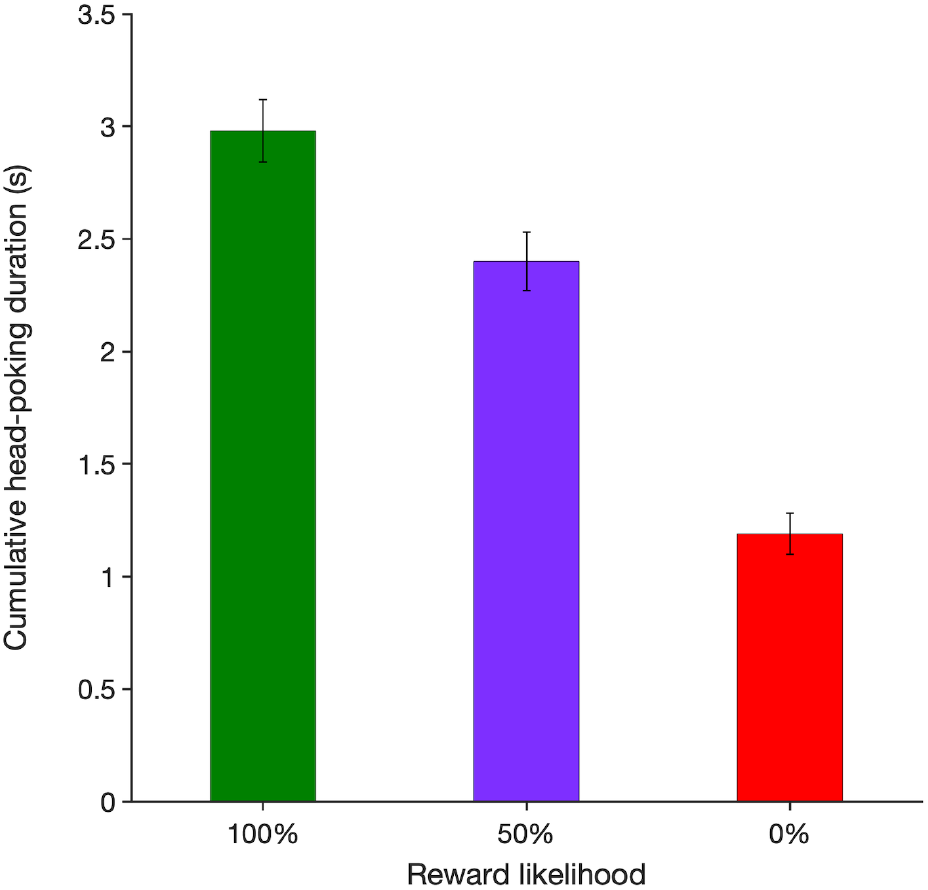
Time spent head-poking into the food magazine during cue presentation in the training phase. The data shows the mean cumulative time (mean ± s.e.m.) subjects spent with their head in the food magazine over the 10s intervals preceding reward outcomes, pooled from the last three sessions. During this time reward-predictive signals indicating a 100% (green), 50% (purple) or 0% (red) chance of reward were presented (corresponding to S^+^, N1 or N2, and S^−^, respectively). n = 24.

### Experiment

#### Preference 1: Choice in 2-option trials

In choice trials a strong preference for *Info* developed in all three treatments, with acquisition occurring more slowly in the *Only_S*^−^ treatment (Fig. 3). A two-way repeated-measures ANOVA with treatment as a between-subject factor, session as a within-subject factor, and (transformed) proportion of choices for *Info* as the response variable, revealed significant effects of treatment (F_2,21_ = 4.00, *P* < 0.05) and session (F_13,276_ = 14.4, *P* < 0.0001), and a significant interaction (F_26,273_ = 1.56, *P* < 0.05), reflecting the slower acquisition in the *Only_S*^−^ treatment. Given the significant interaction, and the plot in Fig. 3, it is obvious that the main effects are caused by rate of acquisition and not by asymptotic levels.

**Fig. 3.**
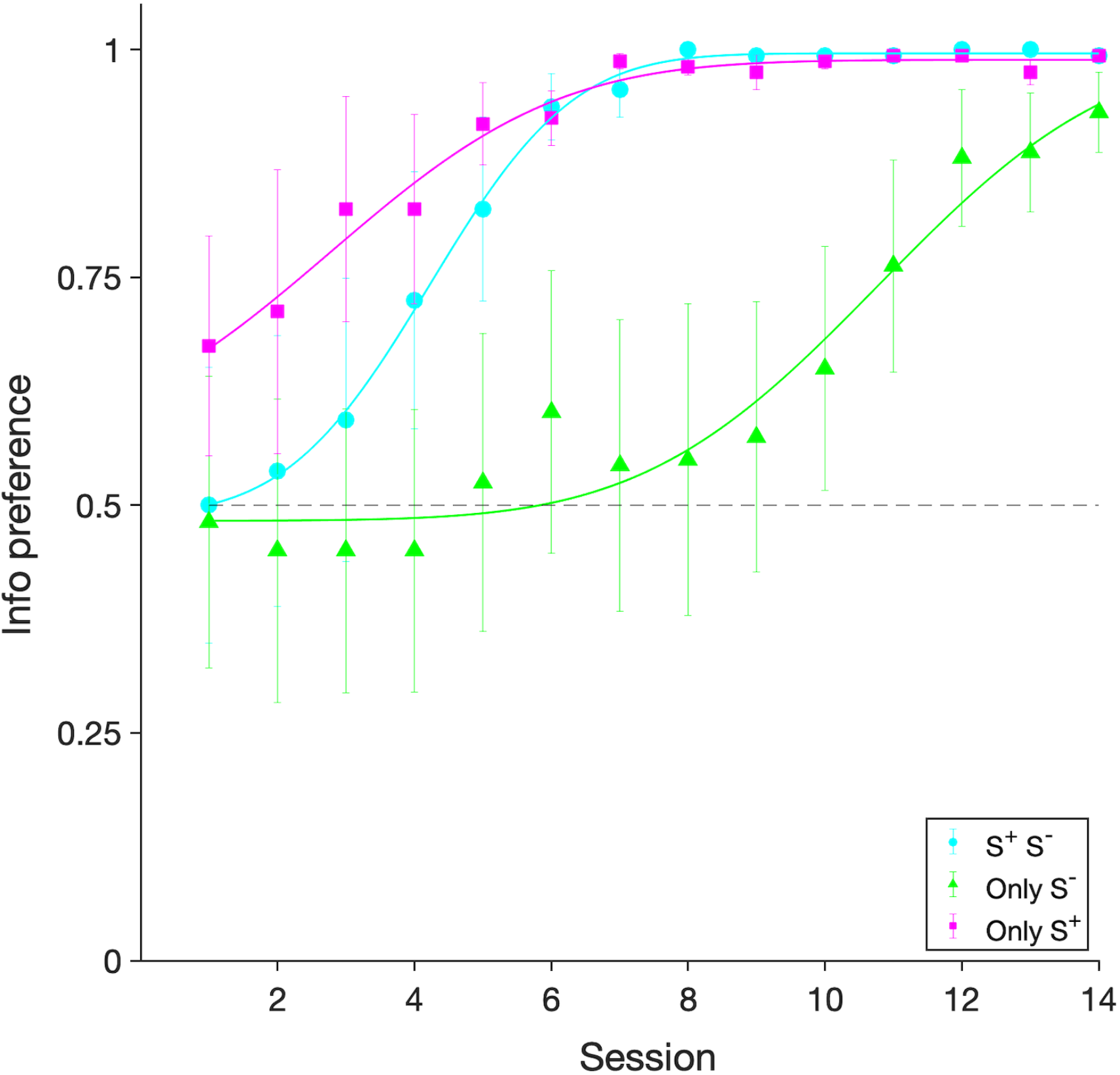
Preference for the *Info* option in choice (2-option) trials. Proportion of observed choices for the *S*^+^_*S*^−^ (n = 8), *Only_S*^−^ (n = 8) and *Only_ S*^+^ (n = 8) groups are shown (means ± s.e.m.) Lines are sigmoidal curves with a cumulative gaussian fit (see methods for details).

We analysed preferences at the end of the experiment by pooling data over the last three sessions. In all three treatments the animals acquired a strong preference for the informative option: 99.8% ± 0.002 (mean± s.e.m.) in the *S*^+^_*S*^−^ treatment; 90% ± 0.03 in *Only_S*^−^, and 98.8% ± 0.005 in *Only_ S*^+^. These values are all significantly greater than 50% (t_7_ = 47.5, *P* < 0.0001; t_7_ = 5.86, *P* < 0.001; and t_7_ = 22.8, *P* < 0.0001, respectively). A one-way ANOVA on proportion of choices pooled from the last three sessions revealed a significant effect of treatment (F_2,69_ = 9.48, *P* < 0.001), with post-hoc pair-wise comparisons showing that bias for *Info* in the *Only_S*^−^ treatment was lower than in the *S*^+^_*S*^−^ treatment (*P* <0.001) and the *Only_S*^+^ treatment (*P* = 0.00459), but was indistinguishable when comparing the *S*^+^_*S*^−^ and *Only_S*^+^ treatments (*P* = 0.68). Again, by reference to Fig. 3 we interpret this as showing that there was a slower rate of acquisition when forthcoming food rewards were signalled by silence than in the other 2 cases, with preference shifting towards a 100% asymptote at different rates in all three treatments.

#### Preference 2: Latency in 1-option trials

While in the previous section we measure preference using proportion of choices in trials when both alternatives were present, here we use latency to respond (reaction time) in single-option forced trials. This is the time between a subject initiating a trial by pressing the back lever and choosing the single option that subsequently becomes available. This has proven to be a robust predictor of choice, and is very informative with respect to the psychological mechanism of choice (see, for instance, Monteiro et al., 2020). Since in each session each individual completed 20 *Info* and 20 *NoInfo* forced trials, we used the median latency shown by each individual for each alternative for analysis. Fig. 4 shows that latencies in single-option trials mirrored the rats’ choice proportions in choice trials: in all treatments latencies were shorter in *Info* than *NoInfo* in the final sessions of the experiment. The absolute value of latencies is shown in Fig. 4 a) and reveals that while latency to respond to *Info* was fairly constant across treatments, latency towards *NoInfo* varied: it was longest in the *S*^+^_*S*^−^, intermediate in *Only_S*^−^ and shortest in *Only_S*^+^. This is interesting because *NoInfo* was identically programmed across treatments. If choices are the result of a horse-race process between the alternatives present at the time of choosing, as claimed under the Sequential Choice Model (Kacelnik et al 2011), then the paradoxical choice phenomenon may be causally mediated by differential behaviour towards the unmodified *NoInfo* option, itself due to manipulations of the *Info* alternative.

**Fig. 4.**
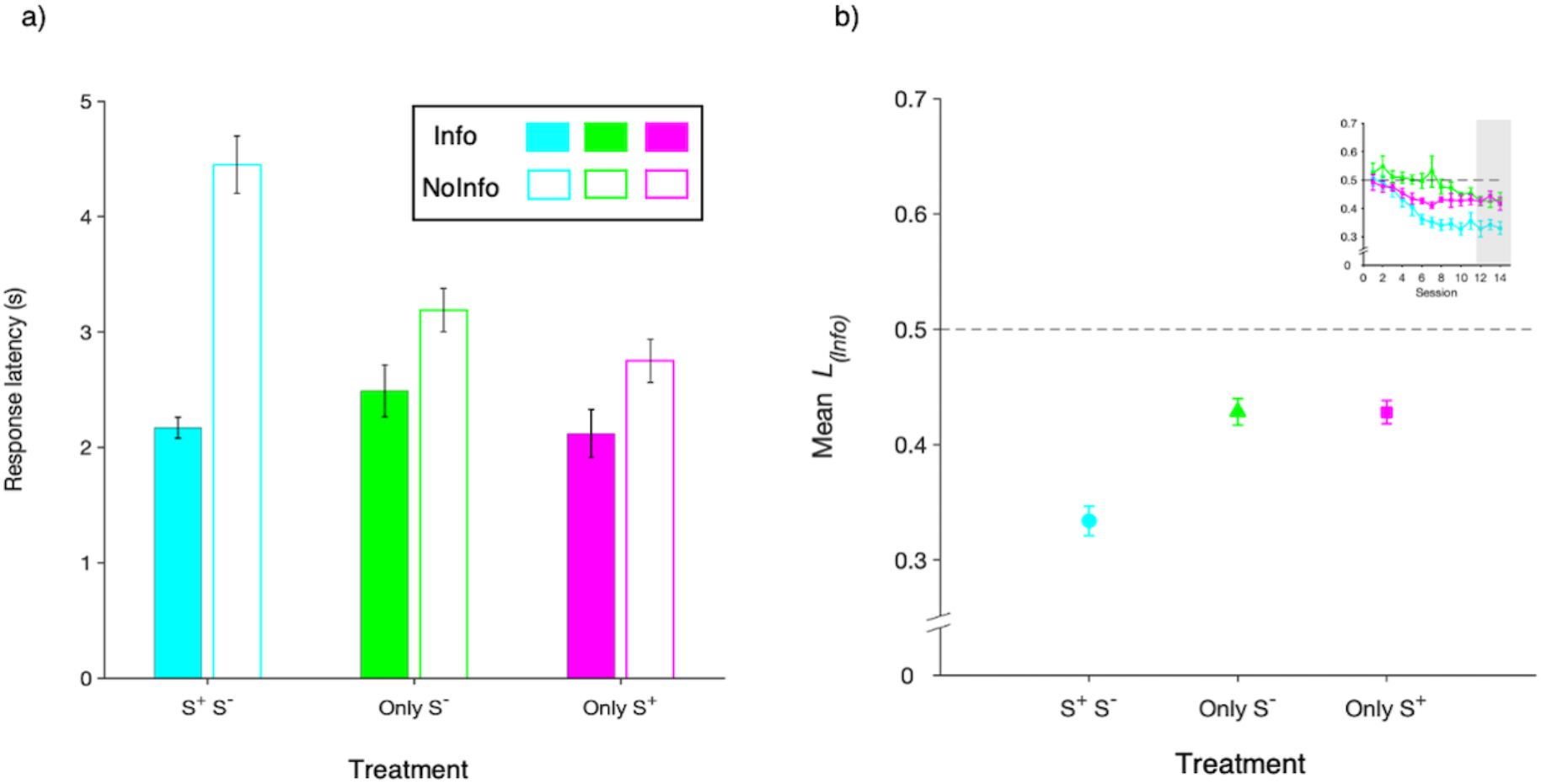
Latency to respond to *Info* vs *NoInfo* in forced (1-option) trials. **a)** Filled bars show latency to respond to *Info* (*R*_*(Info)*_) and unfilled bars show latency to respond to *NoInfo* (*R*_*(NoInfo)*_) across the three treatments (means ± s.e.m.), with data pooled from the last three sessions. n = 8 in each group. **b)** Latency-based preference index as a function of treatment, where *L*_*(Info)*_ = *R*_*(Info)*_ / (*R*_*(Info)*_ + *R*_*(NoInfo)*_), and *R*_*(Info)*_ and *R*_*(NoInfo)*_ are the median latencies to respond in *Info* and *NoInfo* respectively. *L*_*(Info)*_ values below 0.5 indicate preference for *Info* while values of *L*_*(Info)*_ above 0.5 indicate preference for *NoInfo.* The inset shows data across all sessions, with main figure data highlighted in grey.

To quantify preference using forced trials data we ran a two-way repeated-measures ANOVA with treatment as a between-subject factor, session as a within-subject factor and latency-based preference index *L*_*(Info)*_ as the dependent variable (see Methods). This revealed a significant effect of treatment (F_2,21_ = 9.72, *P* < 0.01), session (F_13,273_ = 17.54, *P* < 0.0001), and a significant interaction (F_26,273_ = 2.97, *P* < 0.0001). Post-hoc pair-wise comparisons on data pooled from the last 3 sessions showed that while *L*_*(Info)*_ in *Only_S*^−^ (0.43± 0.012 mean± s.e.m.) and *Only_S*^+^ (0.43± 0.01) were not significantly different from each other (*P = 1*), *L*_*(Info)*_ in both of these groups was significantly higher compared to the *S*^+^_*S*^−^ group (0.33± 0.013; *P* < 0.0001 in both cases). Further, consistently with a preference for *Info*, over the last 3 sessions *L*_*(Info)*_ was significantly lower than 50% in all treatments (*S*^+^_*S*^−^: t_7_ = −7.02, *P* < 0.001; *Only_S^−^:* t_7_ = −3.74 *P* < 0.01; *Only_S*^+^: t_7_ = −4.16, *P* < 0.01), indicating that also on this metric the subjects preferred *Info*.

#### Head-poking during cue presentation

Although behaviour post-choice did not influence outcomes, rats anticipated food by head-poking into the food magazine (possibly a Pavlovian response). Data from choice and forced trials show that in the *Info* option, subjects head-poked more in trials when food delivery was due than when it was not, and showed an intermediate level of head-poking in *NoInfo*, when there was a 50% chance of food delivery (Fig. 5).

**Fig. 5.**
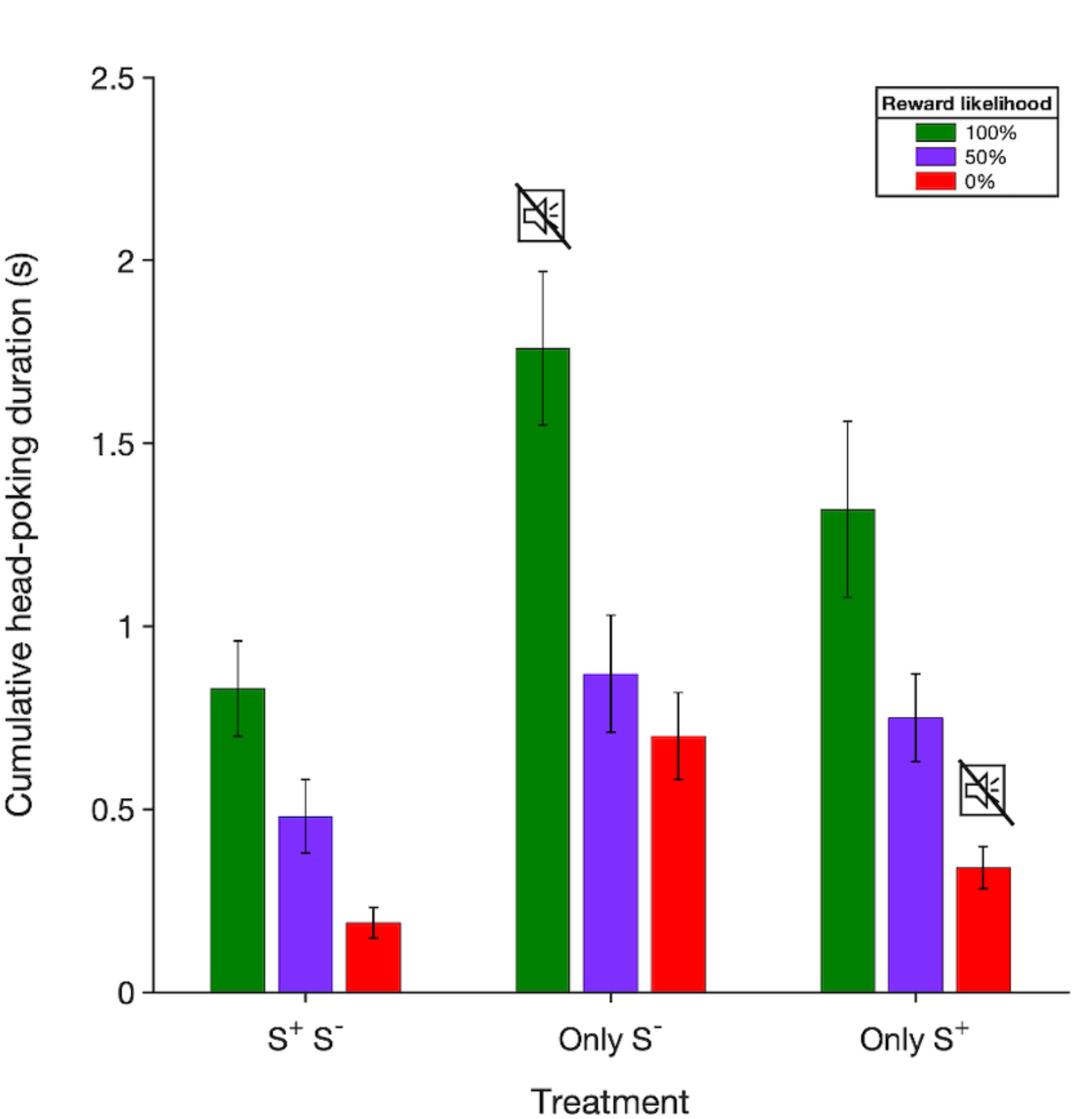
Time spent head-poking into the food magazine between choice and outcome in the main experiment. The bars show the average cumulative time subjects spent with their head in the food magazine in the 10s preceding reward outcomes (± s.e.m.), pooled over the last 3 sessions. During this time reward-predictive signals indicating a 100%, 50% or 0% chance of reward could be presented. Data for the *S*^+^_*S*^−^ (n=8), *Only_S*^−^ (n=8) and *Only_S*^+^ (n= 8) groups are shown. The muted speaker symbol indicates that an explicit cue was not used to signal a particular outcome.

Pooled over the last three sessions, time spent head-poking into the food magazine ranked as reward probability (100% > 50% > 0%). This was the case all treatments: the *S*^+^_*S*^−^ group (100%: 0.83s ± 0.13; mean ± s.e.m., 50%: 0.48s ± 0.1, 0%: 0.19s ± 0.043), *Only_S*^−^ (100%: 1.76s ± 0.21, 50%: 0.87s ± 0.16, 0%: 0.70s ± 0.12), and *Only_S*^+^ (100%: 1.32s ± 0.24, 50%: 0.75s ± 0.12, 0%: 0.34s ± 0.058). A Two-way ANOVA on these data with reward probability as a within-subject factor, treatment as a between-subject factor and cumulative head-poking as the response variable revealed a significant effect of reward probability (100%, 50% or 0% reward; F_13,273_ = 44.38, *P* < 0.0001) but not treatment (F_2,21_ = 2.53, *P* = 0.104). Post-hoc pair-wise comparisons showed that head-poking was significantly higher when reward was due than when it was not in all treatments (highest P < 0.001). The fact that head-poking reflected future reward outcomes differentially in S^+^ and S^−^ trials regardless of treatment shows that rats recognized the contingency they were in regardless of whether an explicit cue was present. Curiously, the absolute level of head-poking seemed to be inversely related to how much food signalling was available (*Only_S*^−^ > *Only_S*^+^ > *S*^+^_*S*^−^), as if attention to explicit signalling competed with exploratory investigation of the food magazine.

## Discussion

We explored the role that two putative psychological mechanisms play in determining preference for an informative option, in which delayed outcomes are signalled by predictive cues, over a non-informative option, in which outcomes remain uncertain until they are realised. A pre-existing observation that is considered to be functionally paradoxical is that in such protocols animals show a strong bias for the informative option, even though the information they gain cannot be used to modify outcomes.

The ‘information hypothesis’, contends that individuals treat uncertainty as aversive, so that informative signals, regardless of whether they bring good or bad news, drive response acquisition. According to this view, both S^+^ and S^−^ should increase preference for the informative option because they reduce uncertainty, regardless of the valence of the outcome with which they are associated. The functional difficulty faced by this hypothesis is that in the experimental situation, reducing uncertainty confers no measurable benefit. This difficulty, like other experimental observations of so-called suboptimal or irrational behaviour, can be addressed post-hoc by arguing that in nature, information about relevant commodities is very often likely to be usable, so that evolution may design utility functions that are somehow tricked by the experimental protocols. Foraging-inspired theoretical models (Freidin and Kacelnik, 2011; Vasconcelos et al., 2015) have argued that in nature, information, even if it announces unfavourable events, is likely to be useful: an animal that knows for sure that the prey being presently pursued will not be captured, would abort the chase, and thus would not pay the opportunity cost of waiting for a null outcome. In other words, it is the artificiality of being unable to use information in the experimental protocol that generates the paradox, which can be reconciled by considering the ecological context in which the mechanism of behaviour evolved (Vasconcelos et al., 2018).

In contrast to the information hypothesis, the ‘conditioned reinforcement’ hypothesis argues that preference for the *Info* option increases due to signals for food (S^+^, ‘good news’) and decreases due to signals for food’s absence (S^−^, ‘bad news’), because S^+^ acquires secondary excitatory properties, and S^−^ inhibitory properties. From this perspective, preference for the ‘informative’ option, where both good and bad news are present, over an option in which neither kind of outcome is signalled (the non-informative option), is a consequence of an imbalance in the quantitative impact of the effect of both signals. Specifically, the excitatory influence of S^+^ is deemed to be greater than the inhibitory effect of S^−^. While mechanistically this is perfectly plausible, this idea also faces functional difficulties. It is not clear *a priori* why the excitatory effect of S^+^ would be greater (or lower) than the inhibitory effect of S^−^. As is often the case, this can also be addressed with reference to the experimental situation being inadequately representative of the environment in which the adaptive learning processes evolved. It is possible that in nature cues indicating the presence of relevant commodities are more prevalent or reliable than those indicating their absence, therefore the power of excitatory and inhibitory conditioned stimuli to modify behaviour need not be symmetric.

We relied on two independent metrics of preference: proportion of choices in 2-option trials, and response latency (reaction time) in 1-option trials. As we show below, this helps to judge the robustness of preferences and to unravel behavioural mechanisms. In the *S*^+^_*S*^−^ treatment, where we reproduced the classic ‘paradoxical choice’ protocol, our results are consistent with previous studies in rats: when presented with two options that differ only in the post-choice predictability of delayed outcomes, rats (as birds and primates) strongly prefer the more informative alternative (Chow et al., 2017; Cunningham and Shahan, 2019; Ojeda et al., 2018). This was observed both in proportion of choices between the alternatives and in differential response latencies when only one of them was present. Furthermore, the absence of either the good or bad news signals in *Info* did not affect asymptotic preferences. The fact that rats preferred *Info* when either only good news or only bad news was signalled indicates that either is sufficient for the acquisition of *Info* preference regardless of the preference metric. This is, prima facie, inconsistent with the conditioning account, because S^−^ is not a likely psychological surrogate of food. However, the relative delayed acquisition when only bad news is present is consistent with the asymmetry between the effects of S^+^ and S^−^ that is part of the conditioned reinforcement hypothesis but not of the information one.

Our analysis of preference on the basis of latency in single option trials is inspired by the Sequential Choice Model (SCM; Kacelnik et al., 2011; Monteiro et al., 2020; Shapiro et al., 2008). The SCM postulates that choice can be modelled as a horserace between the latency distributions of available alternatives, because the alternatives are psychologically processed in parallel, without an active process of choice. Measuring behaviour by more than one procedure is in itself important, because if the phenomenon being measured is meaningful, it should show procedural invariance, a property often claimed to be violated by human studies of choice (Slovic, 1995). We did find consistency between our measures of preference, but also found that using response latency as an additional metric informed about important aspects of potential underlying mechanisms. As Fig. 4 shows, while reaction times in forced trials for *Info* were consistently shorter than in trials for *NoInfo*, across treatments, variations between treatments were mediated only by latency differences in *NoInfo*, which was identically programmed in all three treatments. In other words, treatment effects were mediated by modifications of latency to respond to the least preferred and more constant alternative. This result is striking, could not have been anticipated by the choice results, and is consistent with what was reported by Smith et al. (2018) in a midsession reversal protocol with pigeons, a very different experiment and species. They too, found that changes in choice proportions were explained by variations in latency towards the less preferred alternative in single option trials, when that option did not itself change in its properties. It seems appropriate to infer that parallel processing of alternatives, and mediation through latency variation in less preferred alternatives are very general properties of choice behaviour, something that the most prevalent analysis of choice could not have revealed.

The ‘paradoxical’ preference for unusable information conveyed exclusively through bad news (*Only_S*^−^ treatment) is consistent with observations in starlings (Vasconcelos et al., 2015), monkeys (Lieberman, 1972) and humans (Fantino and Silberberg, 2010; Lieberman et al., 1997), and is difficult to account for exclusively in terms of the conditioned reinforcement hypothesis. The conditioned reinforcement account proposes that through its consistent pairing with food, S^+^ becomes excitatory and S^−^ inhibitory, so that both acquire secondary reinforcing properties of different sign and strength. According to this account, animals prefer the informative option because of the excess excitatory effect of good news. Explanations of precisely how S^+^ can acquire value as a conditioned reinforcer have been developed by different authors and include: the Contrast Hypothesis (Case and Zentall, 2018; Gipson et al., 2009; Zentall, 2013; see also González et al., 2020 for a hypothesis that considers contrast but not conditioned reinforcement *per se*), the Stimulus Value Hypothesis (Smith et al., 2016; Zentall et al., 2015; Smith and Zentall, 2016), the Signals for Good News (SiGN) Hypothesis (Dunn and Spetch, 1990; McDevitt et al., 2016), the Temporal Information Model (Cunningham and Shahan, 2018 though note that their model also considers how primary reinforcement affects choice), and the Selective Engagement Hypothesis (Beierholm and Dayan, 2010; Dinsmoor, 1983).

We do not have the scope or the data to examine and differentiate all these hypotheses in detail, but they all share the assumptions that (1) S^+^ alone is responsible for the acquisition and maintenance of *Info* preference, and (2) The excitatory effect of S^+^ is greater than the inhibitory effect of S^−^ (with some elaborations assuming S^−^ has no effect at all). Were this to be the case, in protocols where S^+^ is substituted by a period without any salient signal, *Info* preference should not be acquired. Our results contradict this prediction of conditioned reinforcement accounts.

It could be argued that the manipulation we performed was not sufficient to eliminate the putative positive conditioned reinforcement afforded by the informative option. After all, head poking data showed that during the post-choice delay rats could anticipate whether or not food was imminent, even for outcomes not signalled by a salient cue (Fig. 5). A conditioning explanation for this could be that subjects treat the compound of their action (lever pressing) plus the immediate absence of a salient cue as a predictive event or conditioned stimulus (CS) in itself, and the delayed outcome (food or no food) as the unconditioned stimulus (US). Thus, they could learn the pairing *[Press Info+Silence]* **→** *food* in the *Only_S*^−^ treatment, while those in *Only_S*^+^ could learn *[Press Info+Silence]* → *no food*. A Pavlovian version of the same idea is that the CS compound does not comprise the rat’s action, but the lever retraction that follows from it. It is therefore possible that in the *Only_S*^−^ group, *Info* lever pressing/retraction followed by the *absence* of an auditory cue is a compound stimulus used by rats to anticipate reward, in other words, it is a virtual S^+^. Under this rationale, conditioned reinforcement can be present in *Info* even with no salient perceptual cue precedes rewards, and thus could account for the results in our experiment.

However, a simple conditioned reinforcement mechanism, like that mentioned above, is unlikely to fully account for *Info* preference across the treatments in our experiment; results from Chow et al. (2017) and Martinez et al. (2017) suggest that the salience of the stimuli in the informative option does influence their efficacy as conditioned reinforcers. Chow et al. (2017) used a blackout to signal forthcoming no reward in the informative option, and found that rats preferred a leaner non-informative option, while Martínez et al. (2017) used the extension of a lever for the same event. Rats preferred the richer (non-informative) one in the first study and the opposite in the latter, which was interpreted by Martinez and colleagues as evidence that for S^−^ to acquire inhibitory properties, it must be salient. The two studies have several methodological differences in addition to the difference in stimuli salience, especially that the probability of food in *Info* and *NoInfo* were 25% and 50% respectively in Chow et al., and 50% and 75% in Martinez et al., but the contrasting results do surely indicate that the salience of S^−^ is an important factor that modulates its ability to acquire secondary reinforcing properties and that this may also be the case for S^+^. Notice that our experiments were yet different: we used reward probabilities of 50% throughout, sounds as explicit signals, and silence (i.e., no change contingent on action) when the absence of a salient stimulus was sought, obtaining very robust preference for the *Info* option in two independent measures of behaviour.

Though asymptotic preferences across the groups in our experiment do challenge the conditioned reinforcement hypothesis, differences in the rate of acquisition are consistent with it. Fig. 3 shows that in the group in which forthcoming food was not signalled (*Only_S*^−^), *Info* preference took significantly longer to be acquired than in the other two groups, where a salient S^+^ was present. This proves that S^+^ enhances the acquisition of *Info* preference. As for S^−^, although some versions of the conditioned reinforcement hypothesis assume that its inhibitory effect is negligible, our results indicate that S^−^ does have an inhibitory effect, probably dependent on its salience. Previous work has shown that rats are more sensitive than pigeons or starlings to losses of intake due to this preference (Martínez et al., 2017; Trujano et al., 2016; Trujano and Orduña, 2015, Fortes et al., 2016; Laude et al., 2014; Fortes et al., 2017; Vasconcelos et al., 2015 and see McDevitt et al., 1997; Pisklak et al., 2015; Spetch et al., 1994). Species differences cannot be used to separate the two hypotheses with presently available data, because both could accommodate them, either through the rewarding value of information or by the parameters of secondary reinforcement processes.

Unlike the conditioned reinforcement hypothesis, the information hypothesis posits that animals prefer the informative option because both S^+^ and S^−^ reduce uncertainty, and this reduction, which is absent in *NoInfo*, is what reinforces preference for *Info*. This explanation makes two predictions that distinguish it from the conditioned reinforcement account. The first is that S^−^ on its own should reinforce *Info* preference. Our main result is consistent with this prediction: in the *Only_S*^−^ group where a salient S^+^ was absent, but S^−^ present, rats also acquired a strong preference for *Info,* indicating that S^−^ reinforces *Info* responses, rather than just inhibiting them.

A strong quantitative version of the information hypothesis could argue that S^+^ and S^−^ should reinforce choices for *Info* to the same extent. This prediction can be derived using Shannon’s (1948) mathematical information theory to quantify the amount of information provided by S^+^ and S^−^. The amount of pre-choice uncertainty associated with *Info* and *NoInfo* is given by:

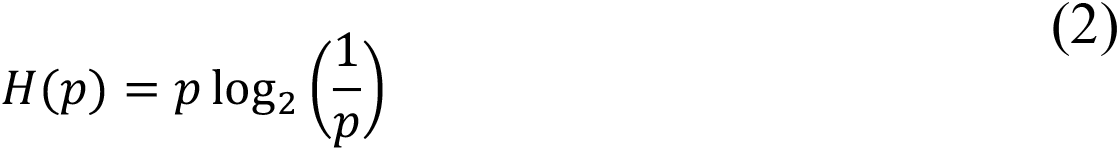

where *H* is the average uncertainty, or entropy of the option, and *p* refers to the probability of the relevant outcome state, which in this case is whether food will occur (Hendry 1969). Since both stimuli in the informative option completely resolve the pre-choice uncertainty, they convey the same amount of information under all possible values of *p*. Thus, if uncertainty reduction were the only consideration, S^+^ and S^−^ would be equally reinforcing. Our finding that its presence/absence modifies rate of acquisition is incongruent with this prediction.

One possibility is that both uncertainty reduction and conditioned reinforcement influence preferences (Daddaoua et al., 2016), so that asymptotic preference is robust to the presence/absence of each stimulus, but rate of acquisition depends on the excitatory/inhibitory role of S^+^ and S^−^ respectively. Our results are consistent with this explanation.

In summary, we have shown that when presented with two equally profitable probabilistic options, rats strongly prefer the one that signals future outcomes over one that does not, even though this information cannot be used to enhance reward intake. Furthermore, this preference is robust to the omission of a salient good news signal and the presence of a salient bad news one in the informative option. We also found that the configuration of signalling contingencies in the informative option affects latency to respond in the non-informative option, and that this may explain the treatment effects on choice. We explored two possible psychological mechanisms: that information is reinforcing *per se* and that cues for good news acquire secondary reinforcing properties that the development of preference. Neither hypothesis on its own can fully account for our findings: our results are consistent with the possibility that both mechanisms operate simultaneously to generate preference. Therefore, both the *amount* of information and its *content* (i.e., good news or bad news) shape the acquisition of preferences in the rat and, possibly, in a range of experimental species.

## Acknowledgements

We’re grateful to Mark Walton for technical advice and useful comments on the manuscript, and technical staff at the BSB for assistance with animal husbandry.

## Declarations

### Funding

This work was supported by funding from the Biotechnology and Biological Sciences Research Council (BBSRC) grant number BB/M011224/1, to VA. AK was sponsored by the Deutsche Forschungsgemeinschaft (DFG, German Research Foundation) under Germany’s Excellence Strategy EXC 2002/1 “Science of Intelligence” –project number 390523135.

### Author contributions

VA, AK, AO and RAM conceptualised and designed the experiment; VA and AO collected the data; VA, AO, TM, and AK analysed the data; VA and AK wrote the first draft of the paper; all authors edited and reviewed the manuscript.

### Competing interests

The authors declare no competing interests.

### Data and materials availability

Data and code for analysis can be made available on reasonable request to the corresponding authors.

## Supplementary figures

**Fig. S1.**
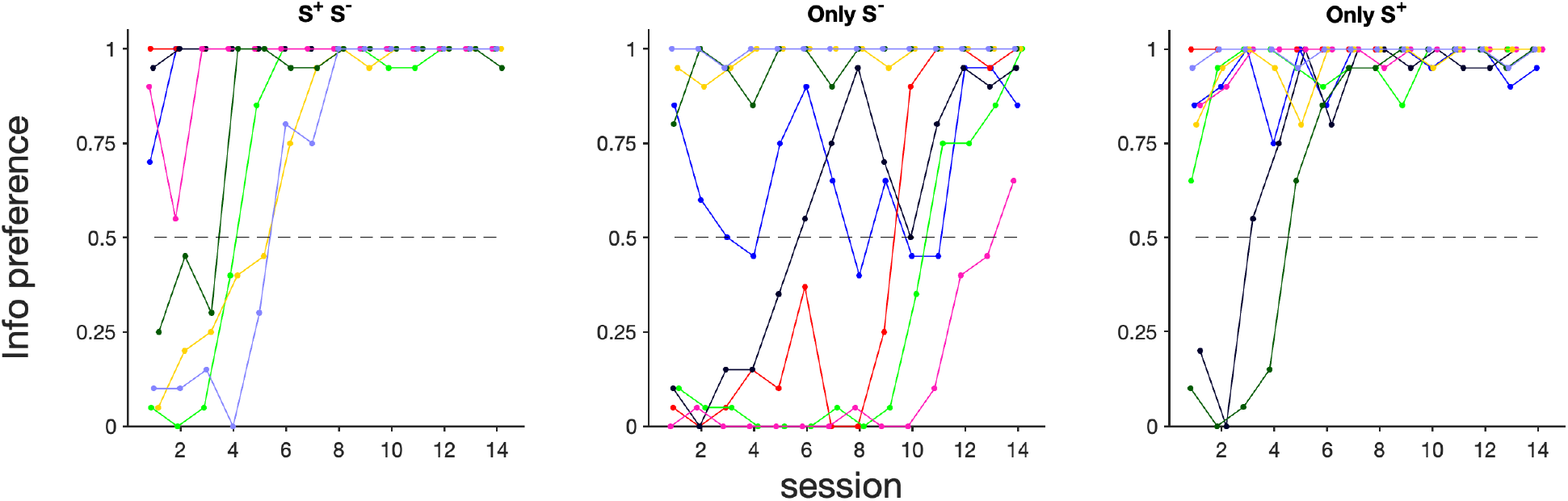
Individual preferences for the *Info* option. Proportion of observed preferences in the *S*^+^_*S*^−^ (n = 8), *Only_S*^−^ (n = 8) and *Only_S*^+^ (n = 8) groups are shown. Each colour represents a different individual for each group.

**Table S1.**
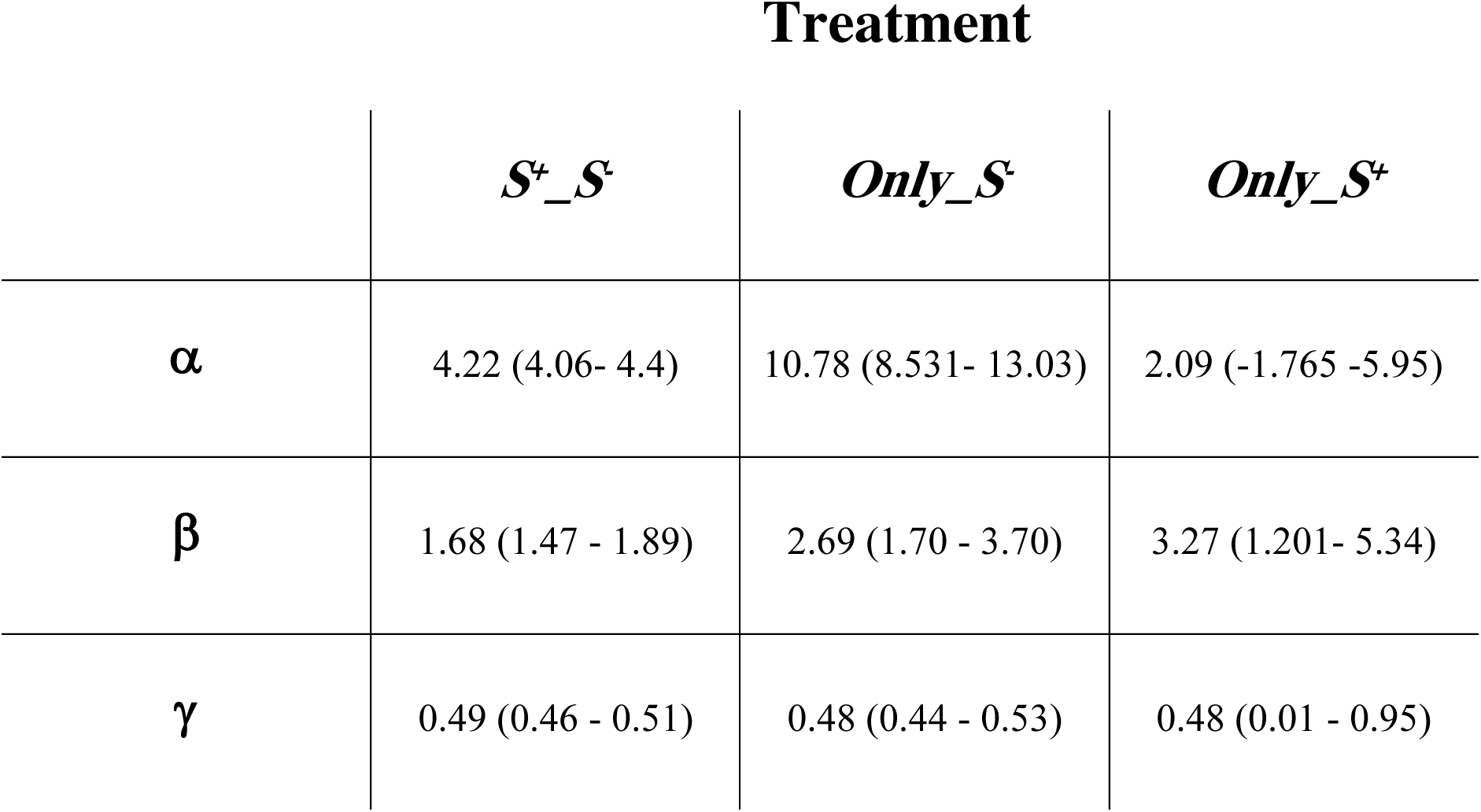
Estimated parameter values from sigmoidal Gaussian curves fit to mean *Info* preference data in the main experiment. *α* gives the inflection point, *β* the slope at the inflection point, and *γ* the lower bound of the curves. The upper bound given by 1-λ was set at 1. 95% confidence bounds are given in brackets.

## Notes

### Competing Interest Statement

The authors have declared no competing interest.

